# PECAM-1 preserves cardiac function in pressure overload-induced biomechanical stress

**DOI:** 10.1101/2021.01.15.426661

**Authors:** Margaret E. McCormick, Mauricio Rojas, John Reader, Ellie Tzima

**Author notes:** **Address for Correspondence:** Ellie Tzima, PhD, Wellcome Centre for Human Genetics, Radcliffe Department of Medicine, University of Oxford OX3 7BN, UK. Heart Failure Discovery Performance Unit, GlaxoSmithKline, 709 Swedeland Road, King of Prussia, PA 19406, USA.

## Abstract

**Background:** Haemodynamic forces play a critical role in proper development of the heart, however much less is known about the mechanisms that regulate cardiac remodelling and function in response to haemodynamic stress in the adult. Platelet endothelial cell adhesion molecule-1 (PECAM-1) is a cell adhesion and signalling molecule that has important roles in regulation of junctional integrity, transendothelial migration and mechanotransduction in response to fluid shear stress. Our previous work identified a role for PECAM-1 in regulating baseline cardiac function via regulation of endothelial-cardiomyocyte communication.

**Methods:** This study investigates the role of PECAM-1 in cardiac remodelling in response to biomechanical stress due to pressure overload induced by transaortic constriction (TAC).

**Results:** Our data reveal that loss of PECAM-1 is associated with systolic dysfunction that is further accentuated following TAC. Adaptive increases in cardiomyocyte cross-sectional area, capillary density and hypertrophic gene expression were all affected with loss of PECAM-1. In control mice, maintained cardiac function was associated with activation of the c-Jun NH(2)-terminal kinase (JNK) pathway, whereas PECAM-1 deletion significantly decreased JNK activation after pressure overload. Our data suggest that in the absence of PECAM-1 signalling, inadequate remodelling of the heart under increased mechanical strain leads to further deterioration of cardiac function, characterized by reduced cardiomyocyte hypertrophy, capillary density and defects in the JNK signalling pathway.

**Conclusions:** Our study reveals a role for PECAM-1 in preservation of cardiac function in response to biomechanical stress induced by pressure overload.

## Introduction

Wall shear stress, the frictional force due to blood flow, is a key epigenetic factor in cardiovascular biology. Initiation of wall shear stress occurs as soon as the heart starts beating early in embryogenesis and plays a crucial role in the proper development of all components of the cardiovascular system (1). It does so by regulating processes critical for embryonic development, including haematopoiesis (2), angiogenesis (3, 4), lymphangiogenesis (5, 6) and remodelling of vessels (7). Specifically, for cardiac development and function, a plethora of studies have demonstrated that shear stress plays an integral role in cardiac morphogenesis and formation of the conduction system (8, 9). However, less is known about the role of haemodynamics in maintaining cardiac homeostasis in the adult.

Given that shear stress plays a key role in regulating endothelial form and function, a significant amount of effort has been dedicated towards identifying the mechanisms by which ECs sense and respond to shear stress. Using various approaches that include knockout cell lines and animals, a number of mechanosensors have been identified (10). These include, but are not limited to, integrins, ion channels, glycocalyx, G proteins and amongst others. Our own work has shown that the junctional cell adhesion molecule PECAM-1 functions as a mechanosensory protein in ECs and confers the ability to sense and respond to the haemodynamic force of flowing blood (11). PECAM-1 is expressed in ECs and haematopoietic cells (12), but not cardiomyocytes (13-15). Although PECAM-1 initiates mechanosignaling specifically in ECs, its presence (or absence) can have profound consequences in the signals transmitted to other cell types, including vascular smooth muscle cells (VSMCs), and thus affect the physiology and pathology of the vessel as a whole. The PECAM-1^-/-^ mouse is viable but displays vascular defects in response to changes in haemodynamic forces. Specifically, PECAM-1^-/-^ mice display impaired flow-mediated dilation due to mis-regulated NO production (16) and reduced VSMC relaxation (17, 18). In addition, our lab has shown that PECAM-1 deficiency results in impaired flow-mediated vascular remodelling that results in reduced VSMC activation and signalling (19). More recently, our laboratory has identified a role for PECAM-1 in the regulation of cardiac contractility and function, despite its lack of expression in cardiomyocytes (15). We showed that PECAM-1 expressed in ECs regulates cardiomyocyte function, as aberrant release of neuregulin-1 (NRG-1) from PECAM-1 KO ECs in the heart results in increased ErbB2 phosphorylation in the cardiomyocytes and dysregulation of contractility.

Given the established role of PECAM-1 in mechanotransduction of haemodynamic stress and its newly identified role in the regulation of cardiac function via endothelial-cardiomyocyte communication, we hypothesized that PECAM-1 is required for the heart to respond effectively to stressed conditions, for instance, compensatory remodelling and hypertrophy in response to pressure overload.

## Materials and Methods

### Animals

PECAM-1^-/-^ C57BL/6 mice were kindly provided by Drs P. and D. Newman (Blood Research Institute, Blood Center of Wisconsin, Milwaukee), bred in-house and used in accordance with the guideline of the National Institute of Health and for the care and use of laboratory animals (approved by the Institutional Animal Care and Use Committees of the University of North Carolina at Chapel Hill). All animal experiments were also approved and authorized by both the University of Oxford Local Animals Ethics and Welfare Committee and by the Home Office, UK. The project licence used in this work was P9133D191. Male (8-10 weeks old) PECAM-1^-/-^ and age-matched, male wild-type PECAM-1^+/+^ littermates were used for all studies.

### Genotyping

Genotyping was determined by PCR analysis of DNA in ear notches, collected for identifying the animals, using the Phire Tissue Kit (F140-WH, Thermo Scientific).

### Transverse aortic constriction

8-10 week old male wild-type (WT) and PECAM-1^-/-^ mice were used for either sham operation or pressure-overload induced by TAC, as previously described (20, 21). Heart function was measured by echocardiography at 7 and 28 days after surgery.

### Echocardiography measurements

Conscious echocardiography was performed as previously described using a Vevo 2100 ultrasound biomicroscopy system (21, 22). All LV dimension data are presented as the average of at least 3 independent waveforms.

### RNA isolation and quantitative PCR

After perfusion with RNA-later (Ambion) under deep anaesthesia, total RNA was isolated from mouse left ventricles using the Qiagen AllPrep Kit (Qiagen, Valencia, CA) following the manufacturer’s protocol. First-strand cDNA was transcribed using random primers and SuperScript II Reverse Transcriptase (Invitrogen). Real-time quantitative PCR was performed using ABsolute SYBR Green ROX mix (Thermo Scientific). Primers used are found in Table 1.

**Table 1.**
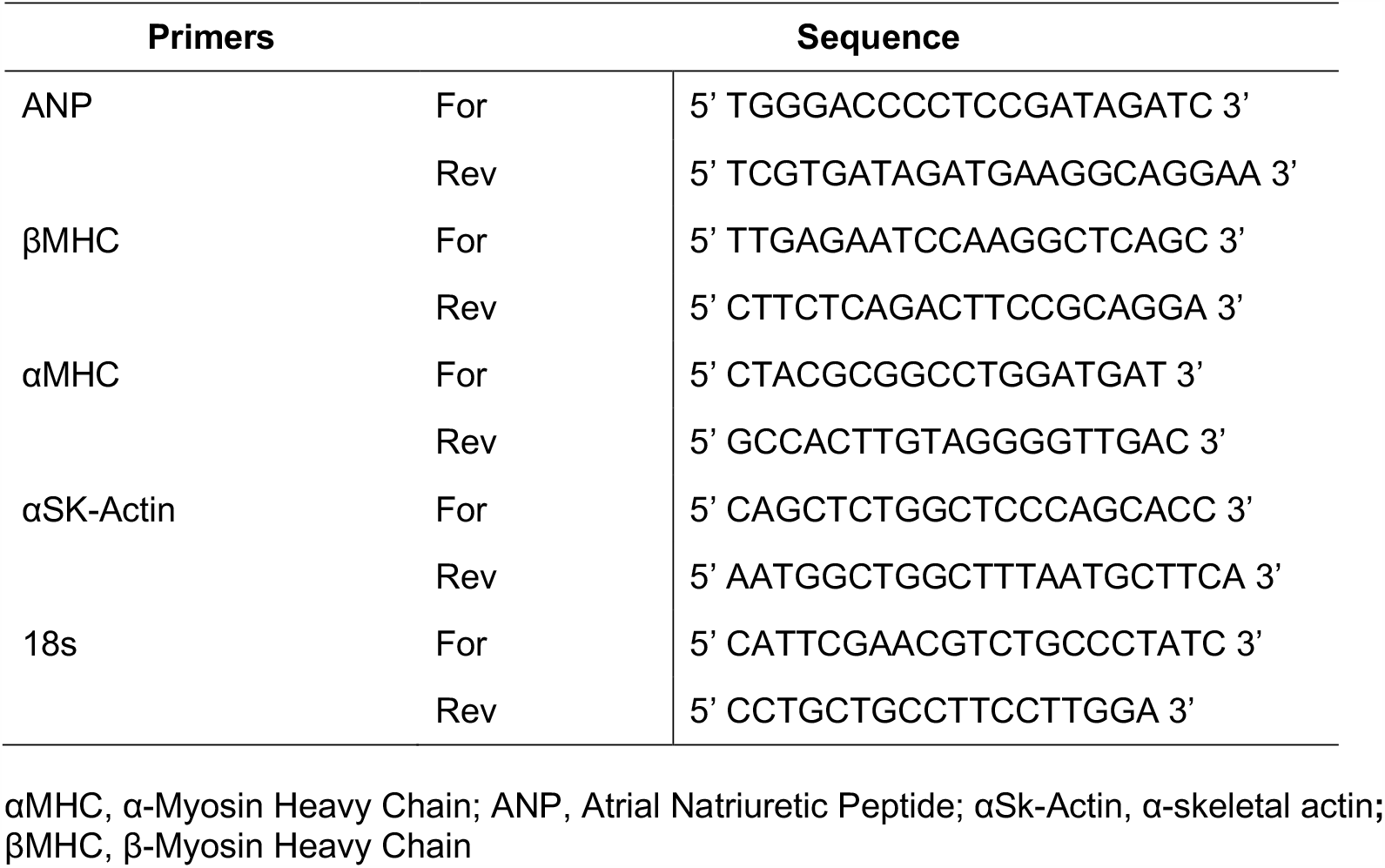
Primers used for quantitative PCR.

### Preparation of lysates and immunoblot analysis

Hearts were homogenized and protein was extracted in a RIPA lysis buffer (50 mM HEPES, 150 mM NaCl, 2 mM EDTA, 0.5% Triton X-100, 0.5% Na-Deoxycholate, 1% NP-40, 2 5mM β-glycerophosphate, 10% glycerol, 1 mM sodium orthovanadate, 1 mM phenylsulphfonyl fluoride, 10 μg/ml leupeptin, 10 μg/ml aprotinin, 10 mM sodium fluoride, 1 mM sodium pyrophosphate, pH 7.2). Protein extracts (30 μg) were subjected to western blot analysis with antibodies against c-Jun NH(2)-terminal kinase (JNK) and phospho-JNK (Cell Signaling). Immunocomplexes were detected using the Licor Odyssey secondary detection system.

### Histological analysis and immunohistochemistry

Hearts were perfused with 4% paraformaldehyde for 24 hours and then switched to 70% ethanol. The hearts were then paraffin-embedded and sectioned into 5-μm sections and stained with haematoxylin and eosin to assess overall morphology ^22^. For cross-sectional analysis of cardiomyocytes and determination of capillary density, heart sections were stained with TRITC-conjugated lectin (*Triticum vulgaris*) and examined by fluorescence microscopy as previously described (21, 22).

### Quantitation and statistical analysis

The band intensity of immunoblots was quantitated using NIH <u>ImageJ</u> software. Each experimental group was analysed using single factor analysis of variance (Excel; Microsoft). Probability values were obtained by performing a 2-tailed Student *t*-test using the same program. Statistical significance was defined as *P<*0.05.

## Results

### Biomechanical stress induces increased cardiac dysfunction in PECAM-1^**-/-**^ **mice**

Previous work from our laboratory demonstrated that absence of PECAM-1 results in impaired cardiac development that is characterized by systolic and diastolic dysfunction (15). To explore the role of PECAM in regulating cardiac remodelling in response to biomechanical stress in the adult, we subjected PECAM-1^-/-^ and PECAM-1^+/+^ littermates to TAC, a well-established model of biomechanical stress and pressure overload. Partial ligation of the aorta leads to increased cardiac haemodynamic stress driving the development of compensatory cardiac hypertrophy(21). Mice were monitored for survival daily, with cardiac function analysed by echocardiography. Following TAC, no differences in survival were observed between PECAM-1^-/-^ and PECAM-1^+/+^ mice. Consistent with previous reports, there were no differences in cardiac dimensions between PECAM-1^+/+^ and PECAM-1^-/-^ sham animals (15). As expected, PECAM-1^+/+^ mice subjected to TAC displayed increased thickness of both the intraventricular septum and anterior wall (Figure 1A, B; Table 2). These anatomical changes were accompanied by preservation of fractional shortening (FS) and ejection fraction (EF) (Figure 2) despite the increase in biomechanical stress. In contrast, 4 weeks after TAC, PECAM-1^-/-^ mice displayed thinner anterior and posterior left ventricular wall thickness and a further increase in LV chamber size (Table 2). Additionally, PECAM-1^-/-^ mice had a significant deterioration in cardiac function in response to pressure overload as measured by both ejection fraction (EF) and fractional shortening (FS) (Figure 2).

**Table 2.**
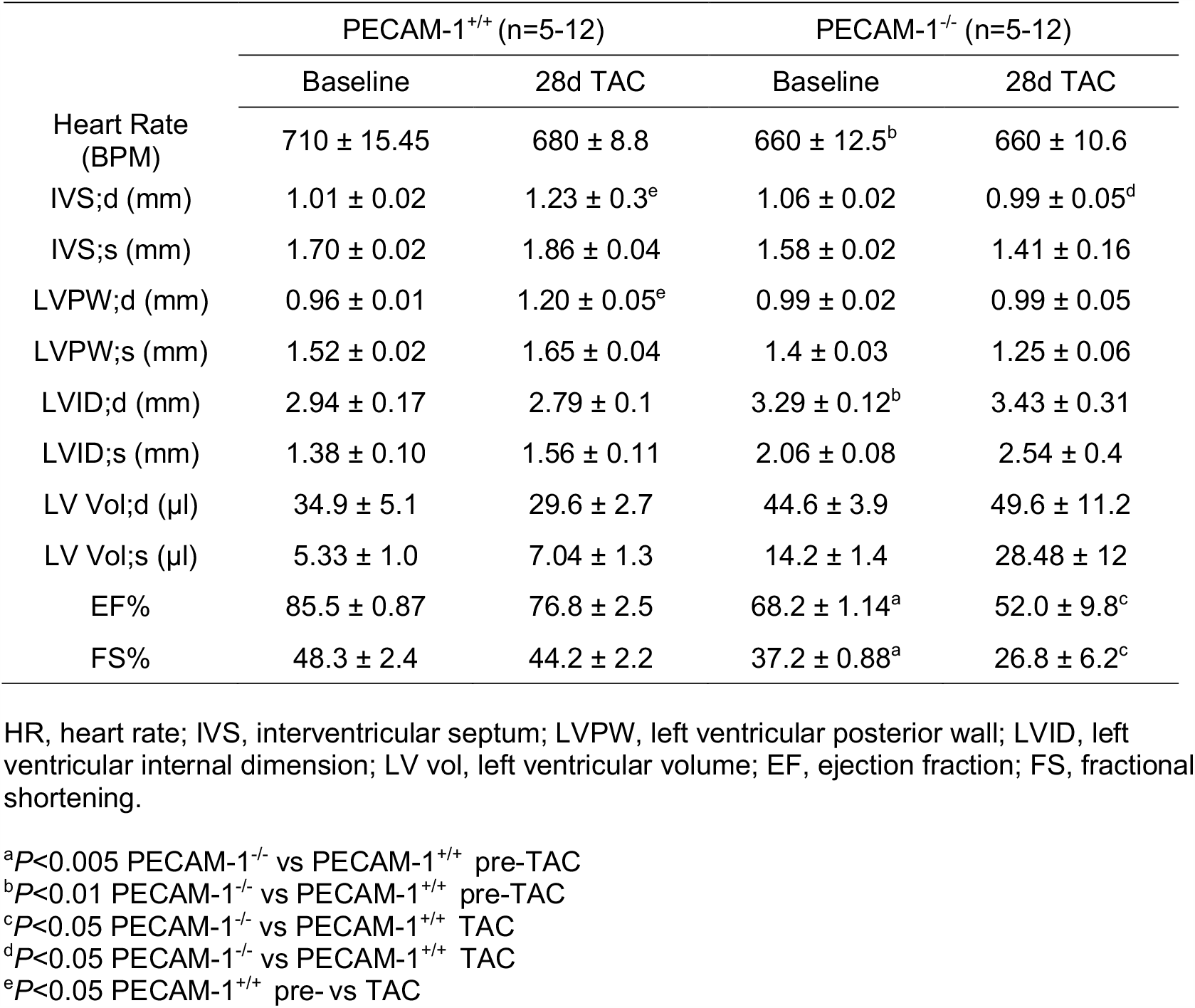
Echocardiographic data from PECAM-1^+/+^ and PECAM-1^-/-^ mice post-TAC.

**Figure 1.**
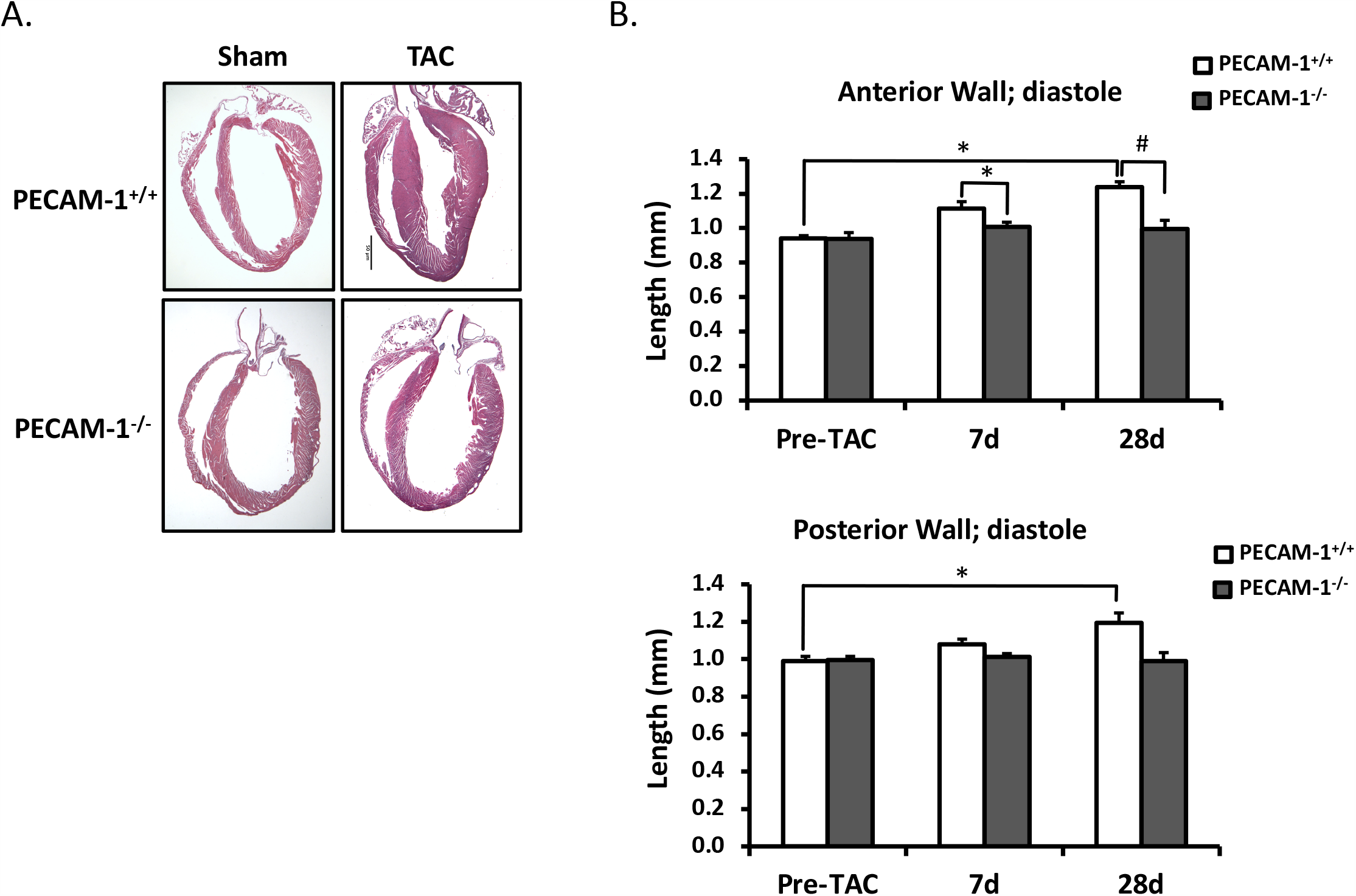
PECAM-1^-/-^ mice display impaired remodelling response to TAC. (A) 8-10week-old PECAM-1^+/+^ and PECAM-1^-/-^ mice underwent either sham or TAC surgeries. Representative H&E stained sections are shown. (B) Left ventricular anterior wall thickness and posterior wall thickness in diastole for sham, 7 day and 28 day PECAM-1^+/+^ and PECAM-1^-/-^ hearts (n=6/genotype for sham, n=10/genotype for TAC, ^*^*P*<0.001 ^#^*P*<0.005). TAC leads to a significant increase in left ventricle anterior and posterior wall thickness in wild-type mice but not in PECAM-1^-/-^ mice.

**Figure 2.**
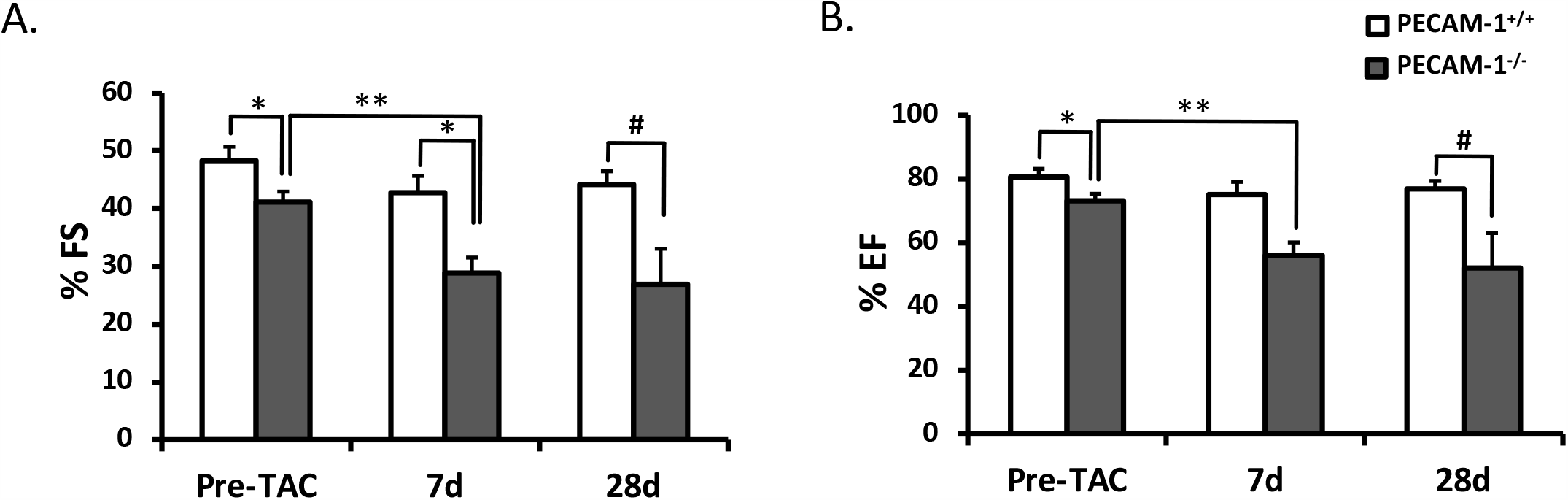
Loss of PECAM-1 induces systolic dysfunction in response to biomechanical stress. (A) Fractional shortening and ejection fraction (B) for sham, 7-day and 28-day hearts (n= 6/genotype for sham, n=10/genotype for TAC, ^*^*P*<0.005 ^**^*P*<0.02, ^#^*P*<0.05). Baseline decreases in FS% and EF% are exaggerated after TAC in PECAM-1^-/-^.

### Reduced hypertrophy and capillary density in the absence of PECAM-1 after TAC

Pressure overload on the heart induces cardiomyocyte hypertrophy characterized by an increase in cardiomyocyte cross-sectional area and wall thickness; this form of hypertrophy is critical for preservation of cardiac function. To assess morphological changes to the heart tissue after TAC, we analysed the cross-sectional area of cardiomyocytes from PECAM-1^+/+^ and PECAM-1^-/-^ hearts (Figure 3A). After 4 weeks of TAC, PECAM-1^+/+^ mice showed a significant increase in cardiomyocyte area, whereas the hypertrophic response was impaired in the PECAM-1^-/-^ hearts (316.0 μm^2^ vs. 247.5 μm^2^) (Figure 3B). Taken together, these data show that PECAM-1 deficiency abrogates the hypertrophic response of the heart to pressure overload and leads to heart failure. Although cardiac dilation and dysfunction is commonly accompanied by left ventricular hypertrophy in order to reduce wall stress, such compensatory hypertrophy was attenuated in the absence of PECAM-1.

**Figure 3.**
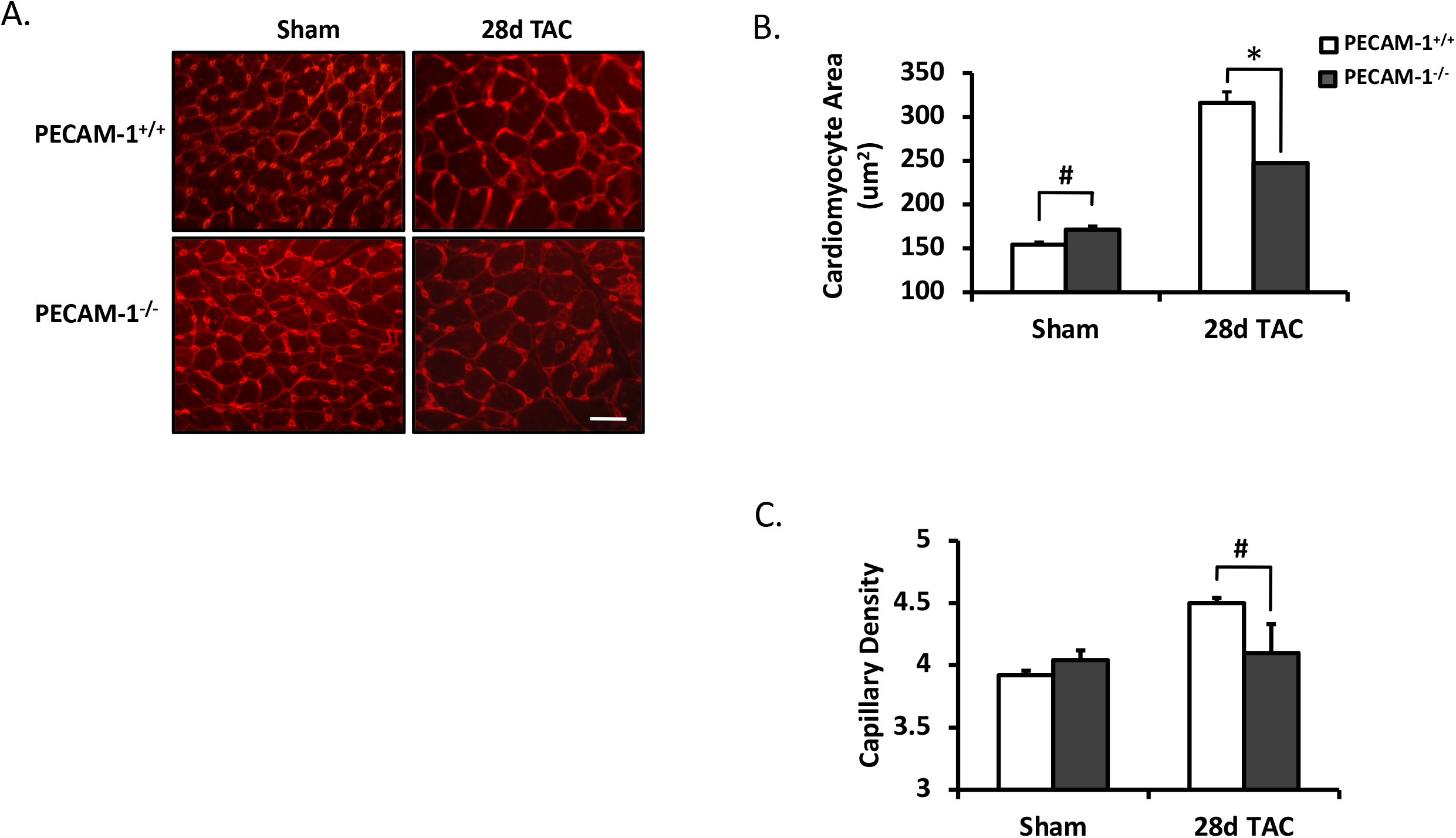
Reduced cardiomyocyte area and capillary density after TAC in PECAM-1^-/-^ mice. (A) TRITC-lectin staining of sham and 28 day TAC PECAM-1^+/+^ and PECAM-1^-/-^ hearts. (B) Quantification of staining in A for baseline and 28 days post-TAC cardiomyocyte cross-sectional area in PECAM-1^+/+^ and PECAM-1^-/-^ mice (n=3-4/genotype, ^*^*P*<0.001, ^#^*P*<0.05). (C) Quantification of staining in (A) for baseline and 28 days post-TAC capillary density in PECAM-1^+/+^ and PECAM-1^-/-^ mice (n=3-4/genotype, ^*^*P*<0.001, ^#^*P*<0.05).

Previous studies have shown that the biomechanical stimulus of pressure overload induces a relatively mild global cardiac ischaemia (23, 24), which is thought to induce angiogenesis and the increased capillary density observed in cardiac hypertrophy (25). We measured capillary density in PECAM-1^+/+^ and PECAM-1^-/-^ mice after TAC and found that whereas PECAM-1^+/+^ hearts displayed increased capillary density, this response was blunted in the absence of PECAM-1 (Figure 3C).

### Impaired cellular activation in PECAM-1^-/-^ hearts after pressure overload

A characteristic of hearts undergoing hypertrophy is the expression of genes that are normally associated with fetal heart development. These include genes that encode protein components of the cardiomyocyte sarcomere and enzymes involved in metabolism, such as upregulation of atrial natriuretic peptide (26), β-myosin heavy chain (βMHC) (27) and skeletal *α*-actin (*α*Ska) (28). The re-activation of the fetal gene programme is considered a protective mechanism for the heart as it switches to glucose as its main energy source in an oxygen-poor environment (29). We measured changes in gene expression after 4 weeks of TAC in the hearts (Figure 4A). Expression of βMHC, ANP, and *α*Ska were increased in both PECAM-1^+/+^ and PECAM-1^-/-^ hearts 7 days after TAC, although this increase was significantly reduced in PECAM-1^-/-^ animals (14-fold in PECAM-1^+/+^ vs. 4-fold in PECAM-1^-/-^ for βMHC). This impaired activation is in agreement with the reduced hypertrophic response seen in the PECAM-1^-/-^ mice (Figure 1).

**Figure 4.**
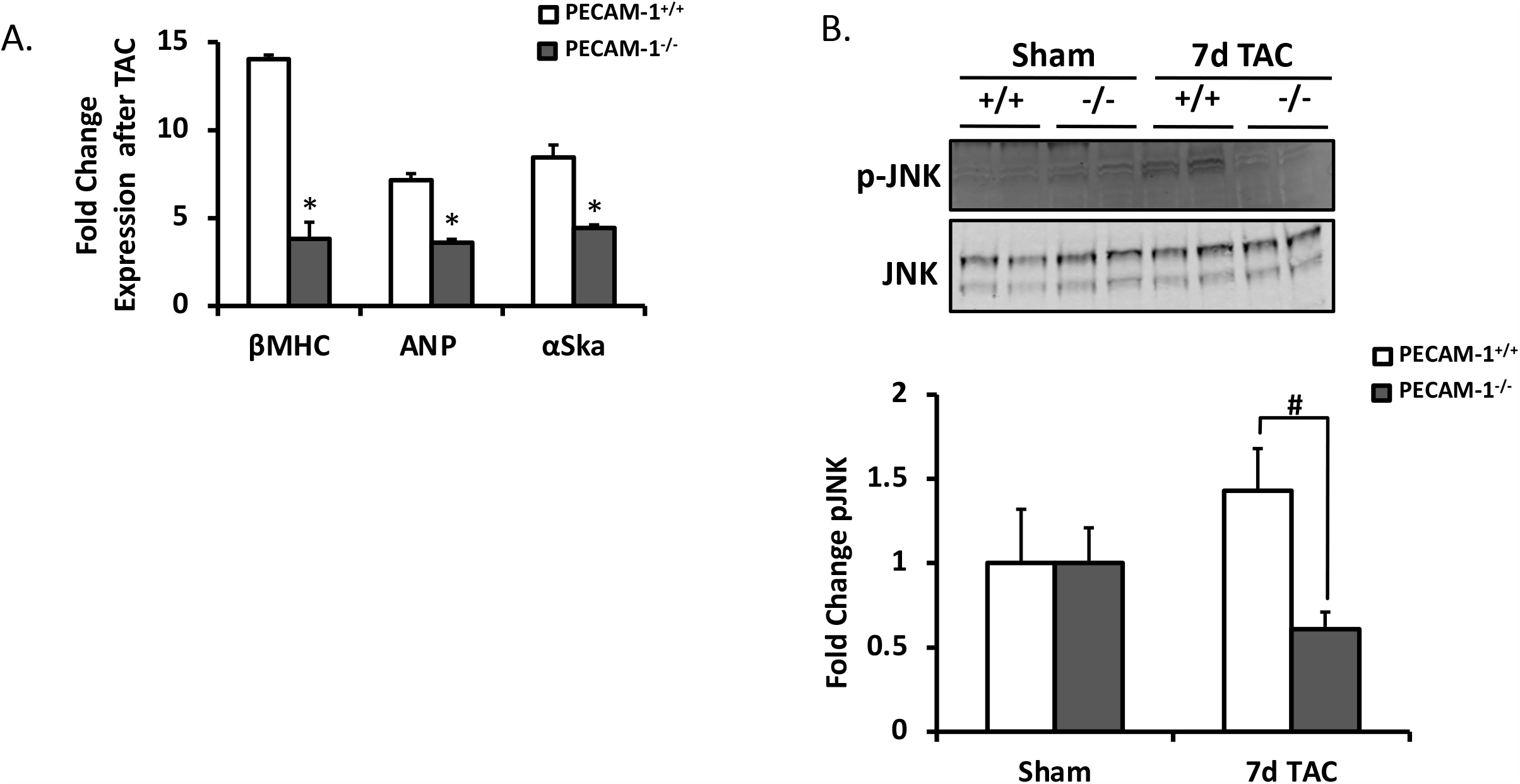
Blunted cellular activation following TAC in PECAM-1^-/-^ mice. (A) Quantitative PCR of cardiac foetal gene expression in 12-week-old male mice. All mRNA species were quantified relative to 18s housekeeping gene expression and presented as fold change (arbitrary units [AU]) relative to sham mice. (n=3/genotype, ^*^*P*<0.01). (B) Representative western blot and quantification for pThr183/Tyr185JNK in sham and 7d post-TAC PECAM-1^+/+^ and PECAM-1^-/-^ hearts. (n=3-4 animals/group) ^#^*P*<0.05

We then investigated molecular mechanisms through which PECAM-1 regulates hypertrophic responses. We examined the JNK pathway, as it has been suggested that JNK plays a protective role in response to pressure overload, protecting the early deterioration in cardiac function following an increase in afterload (30, 31). The JNK pathway has also been shown to be downstream of PECAM-1 in endothelial cells exposed to shear stress (32). JNK phosphorylation at Thr183/Tyr185 was significantly increased after TAC with respect to baseline in control mice, suggesting JNK activation in response to pressure overload. We found that there was significantly less JNK phosphorylation at Thr183/Tyr185 in PECAM-1^-/-^ hearts after TAC, suggesting that PECAM-1 is required for activation of the JNK pathway in response to pressure overload (Figure 4B). Overall, our data reveal a requirement for PECAM-1 in adaptive responses to biomechanical stress imposed due to pressure overload in the heart as loss of PECAM-1 is associated with a reduction in capillary density, cardiomyocyte hypertrophy, reduced activation of the foetal programme, defects in the JNK pathway and, ultimately, deterioration of cardiac function.

## Discussion

The findings presented here demonstrate a requirement for PECAM-1 in regulating the hypertrophic response to biomechanical stress induced by pressure overload. In the absence of PECAM-1, mice fail to remodel and are therefore unable to preserve cardiac function. We also demonstrate that the reduced cardiomyocyte area in PECAM-1^-/-^ hearts in response to biomechanical stress is accompanied by a reduction in capillary remodelling and reduced expression of fetal genes. Mechanistically, these defects are associated with impaired activation of the JNK pathway in the absence of PECAM-1.

Compensatory hypertrophic enlargement constitutes an adaptive response to biomechanical stress; in response to pressure overload, there is an increase in the cross-sectional area of individual cardiomyocytes, which serves to normalise the increase in wall stress induced by mechanical load. This hypertrophy is accompanied by increased capillary density and reactivation of the fetal gene programme. In the absence of external stimuli for enhanced cardiac performance, homeostatic control mechanisms maintain both myocardial size and function in adult hearts. However, increased haemodynamic demands (either due to exercise or increased afterload) lead to cardiomyocyte hypertrophy and an accompanying increase in capillary density(33). A mismatch between myocardial oxygen supply and demand due to lack of an adequate increase in the blood supply can halt adaptation and signal the start of cardiac failure(34, 35). Indeed, Shiojima *et al*. have shown that heart size and cardiac function are dependent on angiogenesis, as disruption of angiogenesis in the heart contributes to the progression from adaptive cardiac hypertrophy to heart failure. (36).

An important observation from our studies is the impaired remodelling in the PECAM-1^-/-^ mice after TAC. Previous studies have demonstrated a requirement for PECAM-1 in remodelling after mechanical stress, particularly responses that require remodelling due to changes in blood flow (18, 19, 37). Specifically, our lab has shown that PECAM-1^-/-^ animals have impaired remodelling in response to changes in haemodynamic stress in models of carotid artery ligation (19) and arteriogenesis (37). In these models, the absence of PECAM-1 prevents the detection of changes in blood flow and subsequent activation of signalling pathways. Based on these studies, we hypothesized that PECAM-1^-/-^ mice would have impaired remodelling after TAC, as TAC dramatically changes the haemodynamic environment within the heart.

It is well-appreciated that intercellular communication in the heart is important not only for proper embryonic development but also for maintaining homeostasis in the adult. Endothelial-cardiomyocyte communication in particular is a two-way street that regulates function of both cell types via secretion of a variety of signalling mediators. It is important to note that although PECAM-1 is expressed in endothelial cells, it is absent from cardiomyocytes (13-15). PECAM-1 is also a well-characterised mechanosensor that is required for sensing of fluid shear stress in endothelial cells(11, 38-41). We therefore hypothesize that the impaired remodelling observed in PECAM-1 KO hearts after TAC is related to the inability of PECAM-1 KO endothelial cells to sense and properly respond to the new haemodynamic environment. Currently, although the role of haemodynamics in cardiac development has been established, much less is known about shear stress in the adult heart. Coronary endothelial cells are mechanosensitive and are thought to utilise the same mechanosensors as those identified in vascular endothelial cells, but this has not been thoroughly investigated(42). Haemodynamics and shear stress are also closely associated with valvular endothelial cell function via changes in gene expression depending on the flow profile experienced (43). Finally, recent studies using 4D flow MRI imaging and next generation sequencing in adult humans and pigs have shown that endocardial endothelial cells are exposed to different shear stresses and display dramatically different phenotypes (44, 45).

It has previously been shown that pressure overload activates the JNK pathway as early as 7 days post-TAC, indicating that JNKs are important in the early response to pressure overload (31). Indeed, JNK1 has been shown to play a critical role in preserving cardiac function in the early phases of haemodynamic stress (31). Importantly, JNK has been shown to be activated by fluid shear stress downstream of the PECAM-1 mechanosensory complex(46). Our data therefore suggest that PECAM-1-dependent JNK activation is associated with pressure overload-induced cardiac remodelling.

In summary, our data suggest that the PECAM-1^-/-^ mouse is a relevant model to study the role of shear stress on adult heart function. When the haemodynamic environment in manipulated (via TAC), PECAM-1^-/-^ animals show impaired remodelling compared to WT animals. Use of this model may help elucidate signalling pathways that are required in shear stress signalling in both cardiac endothelial cells and the heart. A deeper understanding of haemodynamic forces in the heart may reveal novel pharmacologically tractable targets for treating cardiac diseases.

## Data availability

## Acknowledgments

We would like to thank Kirk McNaughton (Histology Research Core Facility, UNC) for help with immunofluorescence and Professor Monte Willis with initial assistance with the echocardiography. We thank the Tzima laboratory colleagues for technical assistance, helpful discussion and critical reading of the manuscript.

